# Targeting the DNM3OS / miR-199a~214 cluster for the treatment of fibroproliferative diseases

**DOI:** 10.1101/242040

**Authors:** G. Savary, M. Buscot, E. Dewaeles, S. Diazzi, N. Nottet, E. Courcot, J. Fassy, K. Lebrigand, I. S. Henaoui, N. Martis, C. Van der Hauwaert, S. Leroy, L. Plantier, A. Paquet, C. L. Lino Cardenas, G. Vassaux, B. Crestani, B. Wallaert, R. Rezzonico, T. Brousseau, F. Glowacki, S. Bellusci, M. Perrais, F. Broly, P. Barbry, C. H. Marquette, C. Cauffiez, B. Mari, N. Pottier

**Affiliations:** Université Côte d’Azur, CNRS, IPMC, FHU-OncoAge, 06560 Valbonne, France; Univ. Lille, EA 4483 - IMPECS - IMPact of Environmental ChemicalS on human health, F-59000 Lille, France; CHU-Nice, Department of Pneumology, FHU-OncoAge, 06000 Nice, France; Centre d’Étude des Pathologies Respiratoires-CEPR, INSERM, UMR1100, Labex Mabimprove, Université François Rabelais, 37000 Tours, France; Assistance Publique-Hôpitaux de Paris, Hôpital Bichat, INSERM U1152, Université Paris Diderot, LABEX Inflamex, DHU FIRE, France; Service de Pneumologie et Immunoallergologie, CHRU Lille, Lille, France; CHU Lille, Service de Biochimie automatisée, Protéines et Biologie Prédictive, F-59000 Lille, France; CHU Lille, Service de Néphrologie, F-59000 Lille, France; Excellence Cluster Cardio-Pulmonary System (ECCPS), German Center for Lung Research (DZL), Justus-Liebig-University Giessen, Giessen, Germany; Univ. Lille, UMR-S 1172 - JPArc - Centre de Recherche Jean-Pierre AUBERT Neurosciences et Cancer, F-59000 Lille, France; CHU Lille, Service de Toxicologie et Génopathies, F-59000 Lille, France

## Abstract

Given the paucity of effective treatments for fibrotic disorders, new insights into the deleterious mechanisms controlling fibroblast activation, the key cell type driving the fibrogenic process, are essential to develop new therapeutic strategies. Here, we identified the long non-coding RNA DNM3OS as a critical downstream effector of TGF-β-induced myofibroblast activation. Mechanistically, DNM3OS regulates this process in *trans* by giving rise to 3 distinct profibrotic mature miRNAs (i.e. miR-199a-5p/3p and miR-214-3p), which influence both SMAD and non-SMAD components of TGF-β signaling in a multifaceted way, through two modes of action consisting of either signal amplification or mediation. Finally, we provide preclinical evidence that interfering with DNM3OS function using distinct strategies not only prevents lung and kidney fibrosis but also improves established lung fibrosis, providing thus a novel paradigm for the treatment of refractory fibrotic diseases such as idiopathic pulmonary fibrosis.

**One Sentence Summary:** The DNM3OS lncRNA is a reservoir of fibromiRs with major functions in fibroblast response to TGF-β and represents a valuable therapeutic target for refractory fibrotic diseases such as idiopathic pulmonary fibrosis (IPF).

## Introduction

Tissue regeneration after injury is a fundamental biological process allowing the ordered replacement of dead or damaged cells to restore tissue integrity and function *(1, 2)*. The current paradigm of wound healing involves two distinct phases: a regenerative phase, in which damaged cells are replaced by cells of the same origin, and a fibrogenic phase, in which scar tissue is generated to protect the injured tissue from further insults until damaged or lost cells are regenerated *(1, 2)*. While the fibrogenic response is usually limited during the normal healing process, in states of iterative or chronic injuries, wound repair can go awry, leading to tissue fibrosis characterized by loss of normal regenerative process, permanent scarring and substantial tissue remodeling *(1, 2)*. The resulting disruption of tissue architecture and organ dysfunction as a consequence of fibrotic changes often leads to severe morbidity and death, especially as the therapeutic options available remain very limited. Indeed, fibroproliferative disorders in aggregate account to as much as 45% of deaths in the industrialized world *(3)* and are increasingly recognized as one of the major forthcoming public health challenges *(4)*.

Myofibroblasts are the key effector cells in the initiation and perpetuation of tissue fibrosis and represent the primary extracellular matrix (ECM) producing cells *(5)*. Myofibroblasts may originate from a variety of cellular sources including resident mesenchymal cells, transdifferentiation of epithelial and endothelial cells, as well as from circulating fibrocytes *(5–7)*. While myofibroblast activation can be triggered by various profibrotic cytokines, research advances over the last two decades have established a prominent role of TGF-β signaling in this process *(8)*. Consequently, intensive efforts are currently devoted towards the discovery and development of drugs able to interfere with this biological pathway for the treatment of fibrotic diseases *(9)*.

Idiopathic Pulmonary Fibrosis (IPF) is a particularly important fibrotic target disease because of its devastating clinical course and the paucity of effective treatment *(10)*. Indeed, nintedanib and pirfenidone are currently the only medication with proven ability to slow disease progression *(11)*. As both of these drugs act primarily on lung fibroblasts by inhibiting essential fibrotic processes mediated by growth factors *(12–14)*, their clinical efficacy provide proof-of-concept that lung fibroblast targeting is a paramount strategy for the development of new anti-fibrotic drugs to cure IPF.

Non-coding RNAs (ncRNAs) comprise multiple classes of RNA transcripts, including microRNAs (miRNAs) and long non-coding RNAs (lncRNAs) that have been shown to exert epigenetic, transcriptional and post-transcriptional regulation of protein coding genes *(15)*. To date, miRNAs are the best characterized ncRNAs, representing a broad class of single-stranded RNAs ~22 nucleotides in length that negatively regulate the stability and/or translation of their target mRNAs *(16)*. While initially presumed to exert important developmental functions, lessons from miRNA loss-of-function studies have instead revealed that these small ncRNAs often profoundly influence stress-induced response of fully-developed tissues, leading to a new paradigm in which miRNA primary function is to maintain homeostasis by buffering stress signaling pathways *(17)*. This, along with the fact that aberrant expression of miRNAs has a causative role in virtually all complex disorders, provide a solid foundation for the rational design of miRNA-based therapies for human diseases with high unmet therapeutic needs *(18)*.

Here, we aimed to comprehensively understand the importance of ncRNAs in the complex molecular events leading to TGF-β-dependent profibrotic activation of lung fibroblasts, and identified a lncRNA named DNM3OS, as a critical TGF-β signaling downstream effector. We further demonstrated that this lncRNA, by serving as a precursor of three distinct fibromiRs (miR-199a-5p, miR-199a-3p and miR-214-3p), is implicated in multiple hallmarks of the profibrotic program orchestrated by lung fibroblasts in response to TGF-β. Finally, we provide preclinical evidence that targeting miR-199a-5p, the most potent profibrotic miRNA of the cluster, may be a promising strategy for the treatment of IPF and other lethal fibrotic diseases.

## Results

### Genomewide profiling of TGF-β-regulated ncRNA in lung fibroblasts

To comprehensively identify ncRNAs that may contribute to lung fibrogenesis, we performed a genomewide assessment of ncRNA expression changes occurring during lung fibroblast activation in response to TGF-β, a major driving cellular event in the development of fibrosis. As expected, RNA-seq analysis of TGF-β-stimulated lung fibroblasts (Datasets 1 and 2) induced large transcriptomic changes (fig. S1) including a TGF-β core signature of 419 coding genes (Fig. 1A and table S1), 130 putative lncRNA candidates (Fig. 1A and table S2) and 46 mature miRNAs (Fig. 1A and table S3). As lncRNA importance has been largely overlooked in fibrogenesis until now, we chose to focus on this class of non-coding transcripts. To identify relevant TGF-β responsive lncRNAs, transcripts were further categorized according to their putative mechanisms of regulation: potential cis-acting lncRNAs (defined as intergenic transcripts whose neighboring gene expression is also modulated by TGF-β or overlapping protein coding gene transcripts whose host gene is also affected by TGF-β) or potential trans-acting lncRNAs (transcripts that does not fall into the aforementioned categories) (fig. S1). Indeed, we reasoned that trans-acting lncRNAs are likely critical intermediates of TGF-β signaling given their propensity to regulate multiple genes at distant locations. We focused our attention on DNM3OS, one of the most TGF-β-induced lncRNAs, transcribed from the negative strand of the dynamin-3 gene (DNM3) (fig. S2, A and B). Conversely, DNM3 transcript is weakly expressed and is not differentially modulated by TGF-β, suggesting that DNM3OS is a genuine trans-acting lncRNA. Upregulation of DNM3OS in lung fibroblasts following TGF-β exposure was independently confirmed by quantitative PCR and RNA FISH analyses (Fig. 1, B and C). In particular, these results showed a strong and early induction of DNM3OS in response to TGF-β (Fig. 1B), which was primarily localized to the nucleus of the fibroblastic cells (Fig. 1C).

**Fig. 1.**
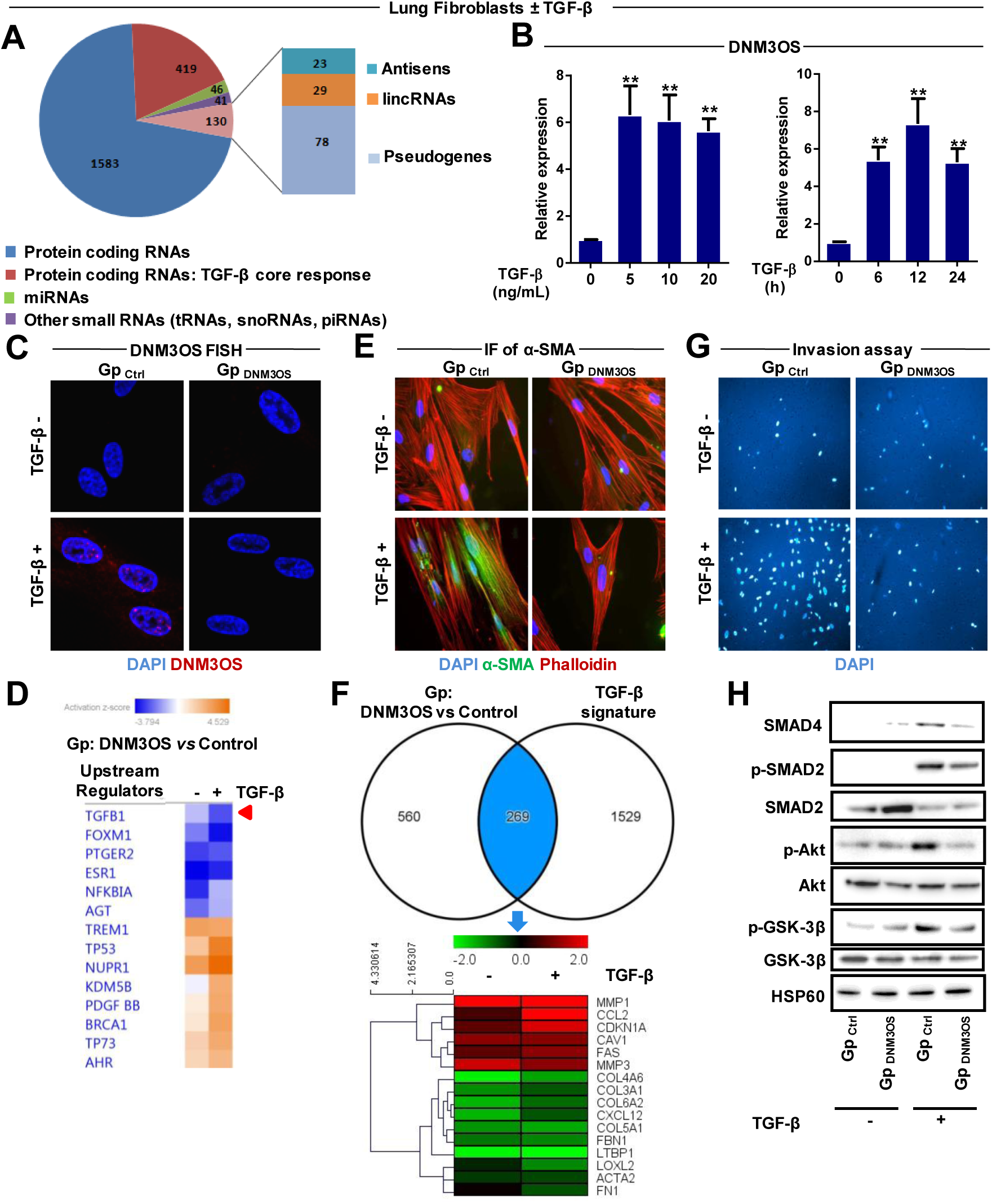
Genomewide profiling of TGF-β-regulated ncRNAs in lung fibroblasts revealed DNM3OS as an essential downstream effector of TGF-β signaling. **(A)** Pie chart illustrating the abundance of the different species of protein-coding and non-coding RNAs differentially modulated by TGF-β as revealed by High Throughput Sequencing (mRNA and small RNA sequencing, Datasets 1 and 2). **(B)** Bar charts showing dose- and time-dependent expression of DNM3OS in MRC5 cells exposed to TGF-β compared to PPIA. Data are expressed as mean ± SEM (*n*=3). ** *P*<0.01. **(C)** RNA-FISH analysis showing the subcellular localization of DNM3OS (red dots) in response to TGF-β using MRC5 cells transfected with either gapmer designed against DNM3OS or gapmer control. Nuclei are stained with DAPI (blue). Representative images out of 2 experiments are shown at x68 magnification. **(D-H)** Functional impact of DNM3OS silencing on TGF-β signaling in pulmonary fibroblasts. MRC5 cells were transfected with either a gapmer designed against DNM3OS or a gapmer control and then incubated with or without TGF-β. **(D)** Heatmap of the predicted upstream regulators after gapmer-mediated silencing of DNM3OS in control or TGF-β conditions (Dataset 3). The red arrow indicates an inhibition of TGF-β-regulated transcripts. **(E)** Immunofluorescence analysis using antibodies against α-SMA (green), phalloidin (red) and DAPI (blue). Representative images out of 2 experiments are shown. **(F)** Venn diagram showing an overlap between genes modulated in response to DNM3OS depletion and an experimental TGF-β signature. Heatmap representing the log2 of the ratio (DNM3OS gapmer/control gapmer) for a subset of 16 typical TGF-β-regulated transcripts **(G)** Invasion assays performed using matrigel. Representative images out of 3 experiments at x10 magnification are shown. **(H)** Western Blot showing both SMAD-dependent (SMAD4, SMAD2, p-SMAD2) and -independent (p-Akt, Akt, p-GSK-3β, GSK-3β) signaling. HSP60 was used as a loading control. One representative experiment out of two is shown.

### DNM3OS is a critical downstream effector of TGF-β signaling in lung fibroblasts

To assess whether DNM3OS influences TGF-β signaling in lung fibroblasts, we performed loss-of-function experiments using gapmers which are defined as single-stranded antisense oligonucleotides (ASOs) that catalyze RNase H-dependent degradation of complementary RNA targets *(19)*. Our results showed that gapmer-mediated silencing of DNM3OS in lung fibroblasts (fig. S2C) strongly affected a subset of genes associated with TGF-β signaling (Fig. 1, D and F and fig. S2D, Dataset 3). Accordingly, we further showed that knockdown of DNM3OS resulted in the inhibition of lung fibroblast differentiation into myofibroblasts (Fig. 1E and fig. S2E), strongly inhibited ECM production (fig. S2E) and lung fibroblast invasion (Fig. 1G and fig. S2F), as well as impaired SMAD and non-SMAD-mediated TGF-β signaling (Fig. 1H). Altogether, these results demonstrate that DNM3OS inhibition abolished the various cellular and molecular phenotypic changes characterizing TGF-β profibrotic effects, highlighting DNM3OS as an essential effector of TGF-β response in lung fibroblasts. Of note, additional genomic analyses indicated that pathways related to actin-based motility, PTEN and Wnt activation were attenuated following DNM3OS inhibition, whereas death receptor signaling and inflammation were likely increased (table S4).

### DNM3OS affects multiple components of the TGF-β pathway by giving rise to 3 distinct profibrotic miRNAs, miR-199a-5p/3p and miR-214-3p

DNM3OS is assumed to function as a precursor of miRNAs as it contains two highly conserved miRNA genes, miR-199a-2 and miR-214 *(20)*. While we previously established miR-199a-5p as a key effector of TGF-β signaling in lung fibroblasts by regulating CAV1 *(21)*, whether the other DNM3OS associated miRNAs also influence TGF-β profibrotic response remains unknown. Our small RNA-Seq data (fig. S1) revealed that, in lung fibroblasts, DNM3OS gives rise to 3 distinct mature miRNAs, namely miR-199a-5p, miR-199a-3p and miR-214-3p (fig. S3A). We then assessed the pulmonary expression of DNM3OS and its associated miRNAs in the well-characterized bleomycin-induced lung fibrosis mouse model *(22)*. Real time PCR analyses were performed on total RNA isolated from the lungs of mice that had been given intratracheal PBS or bleomycin for 2 weeks. In line with our *in vitro* findings, pulmonary expression of DNM3OS and its associated miRNAs was significantly up-regulated during the fibrotic phase of this experimental model (Fig. 2A). Importantly, lungs from patients with IPF also showed increased expression of DNM3OS and its associated miRNAs (Fig. 2B). We thus hypothesized that DNM3OS promotes TGF-β signaling by serving, at least in part, as a precursor of three fibromiRs, which regulate distinct targets involved in this profibrotic signaling cascade. In line with this, our results showed that, in lung fibroblasts, expression of these fibromiRs mirrored that of DNM3OS in response to TGF-β, whereas DNM3OS silencing resulted in a marked reduction of its associated miRNAs (Fig. 2C and fig. S3, A to C). Of particular interest, gain- and loss-of-function approaches showed that miR-214-3p and miR-199a-5p similarly influence lung fibroblast profibrotic response to TGF-β by promoting their differentiation into myofibroblasts (Fig. 2, D and E and fig. S3D) and enhancing their invasive properties (Fig. 2, F and G and fig. S3E). In contrast, modulation of miR-199a-3p expression had no effect on the phenotypic changes occurring during TGF-β-induced activation of lung fibroblasts (Fig. 2, D to G and fig. S3, D and E). Taken together, these results suggest that miR-199a-5p and miR-214-3p likely target functionally related genes within the TGF-β signaling cascade whereas miR-199a-3p is likely implicated in the regulation of a distinct signaling component. To further assess how miR-199a-3p and miR-214-3p influence TGF-β response, we first identified the cellular pathways and gene target candidates regulated by each of these miRNAs using a combination of experimental and *in silico* approaches described earlier *(21, 23, 24)*. Although a specific gene expression pattern was identified for each miRNA overexpressed in lung fibroblasts (fig. S4A, Dataset 4), GSEA algorithm *(25)* revealed a significant association between these miRNA-derived gene expression profiles and an experimental signature of lung fibroblast response to TGF-β (fig. S4B, Dataset 5). Consistent with our *in vitro* findings, this analysis showed a significant overlap between genes down-regulated by TGF-β and those repressed by either miR-199a-5p or miR-214-3p, whereas this link was weaker for miR-199a-3p (fig. S4B). Furthermore, functional annotation of the gene expression profiles associated with miRNA overexpression also retrieved a partial overlap for canonical pathways associated with fibrosis such as “FGF signaling” “HGF signaling” “PI3K signaling”, “PTEN signaling” or “Wnt / β-catenin signaling” (table S5). We next looked for an enrichment of putative direct targets in transcripts that were down-regulated after mimic overexpression (fig. S4, C and D). Of these putative targets, we focused our attention on those associated with the most significant canonical pathways described above (fig. S4E). This uncovered two important negative regulators of TGF-β signaling, GSK3B (coding for GSK-3β) and PTGS2 (coding for COX-2), as putative gene targets of miR-214-3p as well as two potent anti-fibrotic genes, HGF and FGF7 (also known as KGF), as potential targets of miR-199a-3p (fig. S4E).

**Fig. 2.**
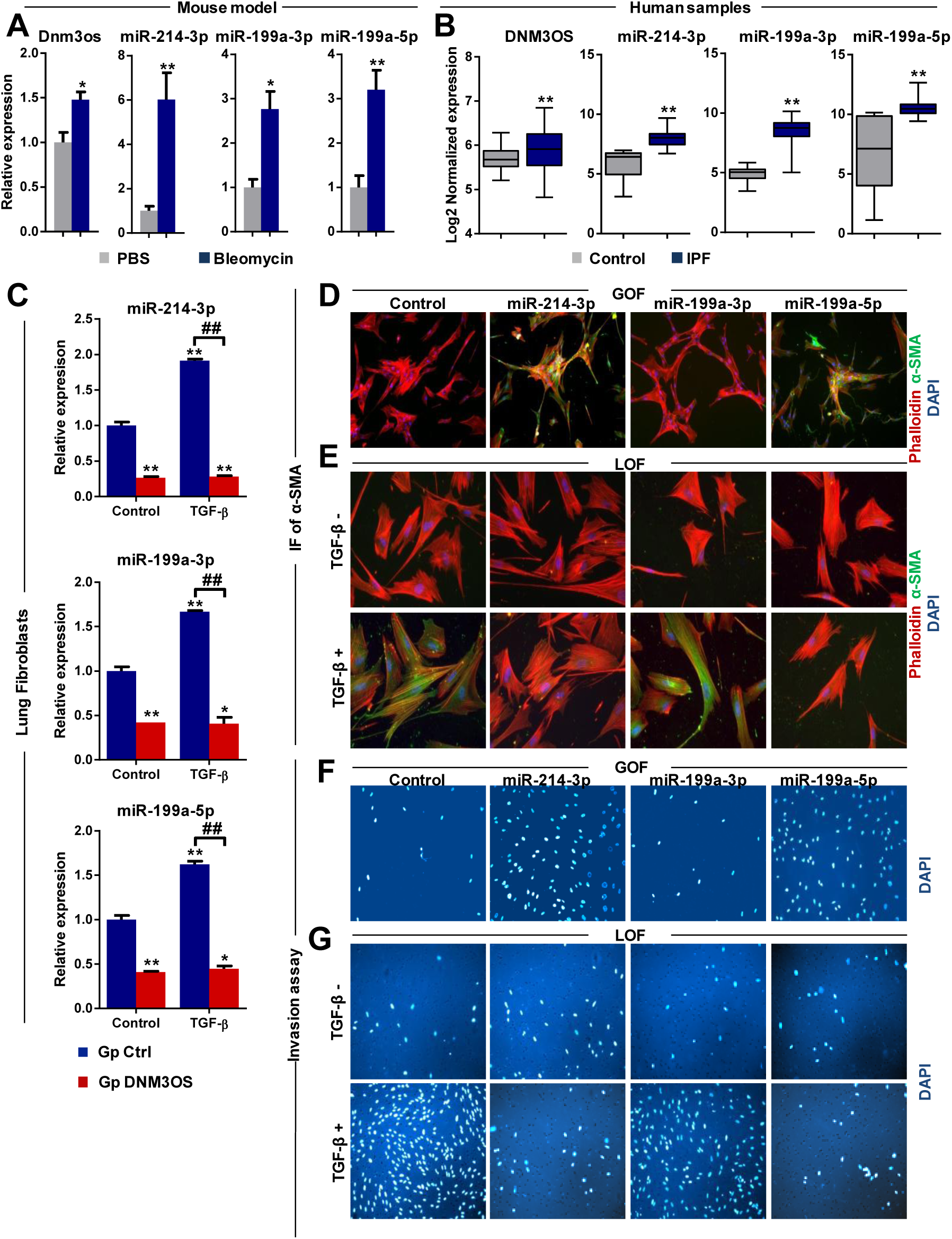
Altered expression of DNM3OS and its associated miRNAs in pulmonary fibrosis and functional impact of DNM3OS-associated miRNAs on lung fibroblasts. **(A)** Bar charts showing the relative pulmonary expression of DNM3OS and its associated miRNAs in C57BL/6 mice treated either with bleomycin or PBS for 14 days. Data are expressed as mean ± SEM (*n*=4-6 in each group). * *P*<0.05, ** *P*<0.01. **(B)** Box plots showing the log2-transformed normalized expression of DNM3OS and its associated miRNAs in both IPF (*n* = 118 for DNM3OS and n=19 for the mature miRNAs) and control (*n* = 49 for DNM3OS and *n*=6 for the mature miRNAs) lungs. The box represents the 25–75% quartiles, the line in the box represents the median and whiskers represent the range. ** *P*<0.01. **(C)** Bar charts showing the relative expression of DNM3OS-associated mature miRNAs compared to RNU44 following DNM3OS silencing. MRC5 cells were transfected with either a DNM3OS gapmer or a control gapmer and then incubated with or without TGF-β. Data are expressed as mean ± SEM. (*n*=3) * *P*<0.05 ** *P*<0.01 ## *P*<0.01. **(D-G)** Gain (GOF) and loss (LOF) of function experiments showing that only miR-214-3p and miR-199a-5p mediate TGF-β-induced lung fibroblast invasion and differentiation into myofibroblasts. MRC5 cells were transfected with pre-miRNA mimics, LNA-based miRNA inhibitors or their respective controls, then incubated with or without TGF-β. **(D,E)** Immunofluorescence analysis using antibodies against α-SMA (green), phalloidin (red) and DAPI (blue). Representative images out of 2 experiments are shown at x20 magnification. **(F,G)** Invasion assays performed using matrigel. Representative images at x10 magnification are shown for each condition.

In other processes than fibrogenesis, PGE2, the enzymatic product of COX-2, and GSK-3β have been functionally linked to β-catenin pathway, a SMAD independent profibrotic component of TGF-β signaling *(26)*. As nuclear β-catenin accumulation in fibroblasts is a critical fibrogenic event *(27)*, we hypothesized that COX-2 and GSK-3β act in concert to hinder TGF-β-induced nuclear β-catenin translocation and that miR-214-3p fine tunes this regulatory loop. Accordingly, our results established both PTGS2 and GSK3B transcripts as *bona fide* targets of miR-214-3p (fig. S5, A and B). We further showed that miR-214-3p ectopic expression in lung fibroblasts repressed GSK-3β (fig. S5C) and was sufficient to induce β-catenin nuclear translocation (Fig. 3, A and B). Importantly, our results also showed that siRNA-mediated depletion of GSK-3β in lung fibroblasts phenocopies miR-214-3p overexpression, with a strong increase in α-SMA expression levels (fig. S5, D and E). Similarly, we also demonstrated that miR-214-3p modulates COX-2 expression and consequently PGE2 secretion (Fig. 3C and fig. S5, F and G). Interestingly, we provide new mechanistic insights into PGE2 anti-fibrotic effect by showing that this prostaglandin specifically inhibits the non-SMAD TGF-β signaling pathway, in particular the GSK-3β/β-catenin component (Fig. 3, D and E). Finally, as IPF fibroblasts exhibit a defective COX-2/PGE2 axis which has been associated with a reduced sensitivity to apoptosis *(28)*, we also showed that miR-214-3p overexpression renders lung fibroblasts more resistant to FASL-mediated apoptosis (fig. S5, H to J).

**Fig. 3.**
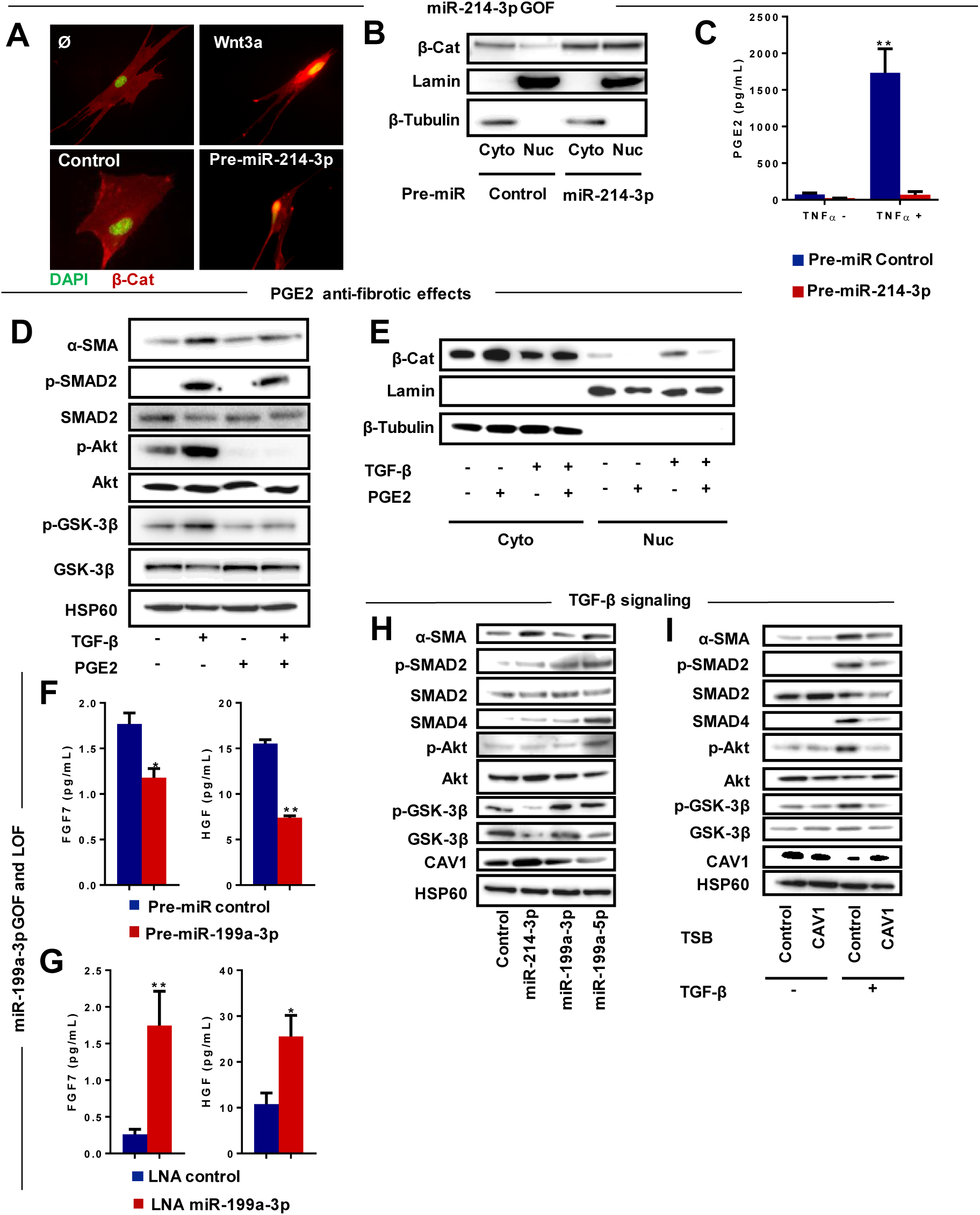
GSK-3β/COX-2 and KGF/HGF are direct targets of miR-214-3p and miR-199a-3p respectively. **(A,E)** Gain of function (GOF) experiments showing that miR-214-3p promotes β-catenin nuclear translocation by targeting GSK-3β and COX-2 in lung fibroblasts. **(A)** Immunofluorescence analysis showing the nuclear translocation of β-catenin using antibody against β-catenin (red) and DAPI (green). MRC5 stimulated with WNT3A were used as positive control. Representative images out of 2 experiments are shown at x40 magnification. **(B)** Western blot analysis of β-catenin levels in cytosol and nuclear fractions. β-tubulin and lamin A/C were used as cytosolic and nuclear loading controls, respectively. **(C)** Gain of function experiments showing that miR-214-3p suppresses PGE2 release from lung fibroblasts by targeting COX-2. **(D,E)** PGE2 hinders TGF-β-induced myofibroblast differentiation by inhibiting β-catenin nuclear translocation. **(D)** Western blot showing the impact of PGE2 on myofibroblast differentiation (α-SMA) as well as both SMAD (p-SMAD2 and SMAD2) and non-SMAD (p-Akt, Akt, p-GSK-3β, GSK-3β) signaling pathways. HSP60 was used as a loading control. One representative experiment out of two is shown. **(E)** Western blot analysis of β-catenin levels in cytosol (cyto) and nuclear (nuc) fractions after either PGE2 or TGF-β exposure or both. β-tubulin and lamin A/C were used as cytosolic and nuclear loading controls, respectively. **(F-G)** Gain (GOF) and loss (LOF) of function experiments showing that miR-199a-3p mediates the suppressive effects of TGF-β on FGF7 and HGF expression in lung fibroblasts. Bar charts showing FGF7 and HGF quantification using ELISA assays. Data are expressed as mean ± SEM (*n*=3). * *P*<0.05 ** *P*<0.01. **(H,I)** miR-199a-5p by targeting CAV1 has a prominent role on TGF-β signaling in lung fibroblasts **(H)** Western Blot showing the impact of individual DNM3OS-associated miRNA overexpression in lung fibroblasts on α-SMA and CAV1 expression as well as on both canonical (p-SMAD2, SMAD2 and SMAD4) and non-canonical (p-Akt, Akt, p-GSK-3β, GSK-3β) TGF-β signaling pathways. HSP60 was used as a loading control. One representative experiment out of two is shown. **(I)** Western Blot showing that specifically inhibiting miR-199a-5p binding on hCAV1 3’UTR using CAV1 TSB abolishes TGF-β-induced pulmonary fibroblast activation by inhibiting both SMAD (p-SMAD2, SMAD2 and SMAD4) and non-SMAD (p-Akt, Akt, p-GSK-3β, GSK-3β) signaling pathways.

HGF and FGF7 are two fibroblast-derived growth factors sharing similar anti-fibrotic activities, in particular the promotion of epithelium repair, and whose expression is inhibited by TGF-β *(29, 30)* (fig. S6A). Although HGF has been previously validated as a direct target of miR-199a-3p in fibroblasts *(30)*, whether FGF7 is also a genuine target of miR-199a-3p was not known. Our results established FGF7 as a direct target of miR-199a-3p (fig. S6B) and further demonstrated by gain- and loss-of-function approaches that this miRNA likely mediates TGF-β-induced downregulation of both FGF7 and HGF in pulmonary fibroblasts (Fig. 3, F and G and fig. S6C). Interestingly, our results strongly suggest that DNM3OS through miR-199a-3p and miR-214-3p fine tunes the previously described HGF/COX2/PGE2 anti-fibrotic axis *(31)*.

Finally, our *in vitro* results underscore a prominent role of miR-199a-5p in the regulation of TGF-β signaling in lung fibroblasts notably by promoting both SMAD and non-SMAD pathways (Fig. 3H). In addition, using a CAV1 Target Site Blocker (TSB), we provide mechanistic evidence that these two pathways are controlled through a miR-199a-5p-CAV1-feedforward regulatory circuit (Fig. 3I). Altogether, our data support a molecular scenario where DNM3OS promotes TGF-β signaling in lung fibroblasts by giving rise to three fibromiRs which collectively target distinct components of the signaling cascade.

### miR-199a-5p promotes lung fibrogenesis in mice through a CAV1-dependent mechanism

Given the relative ease by which miRNAs can be targeted *in vivo (32)*, we decided to further assess the relevance of our findings by focusing on miR-199a-5p using the bleomycin-induced lung fibrosis mouse model (Fig. 4A). Consistent with our *in vitro* findings, pretreatment with *in vivo* LNA–modified antisense probes designed against miR-199a-5p prevented the enhanced pulmonary expression of miR-199a-5p, the inhibition of CAV1 and the induction of ECM proteins, including fibronectin and collagen, in response to bleomycin administration (Fig. 4, B and C and fig. S7A). Moreover, H&E staining demonstrated striking attenuation of bleomycin-induced lung fibrosis in mice pretreated with miR-199a-5p antisense probes (Fig. 4B); this finding was confirmed by sirius red staining, which highlights collagen deposition (Fig. 4B). Likewise, LNA-mediated silencing of miR-199a-5p prevented the accumulation of myofibroblasts in bleomycin-treated lungs, as demonstrated by immunohistochemistry staining with anti α-SMA antibody (Fig. 4B). These results were also confirmed at the transcriptome level, by showing a strong attenuation of the vast majority of bleomycin-regulated transcripts in mice treated with miR-199a-5p inhibitor (Fig. 4D and fig. S7B, Dataset 6). Remarkably, similar results were obtained *in vivo* using Cav1 Target Site Blockers (Fig. 4A and fig. S7C, Dataset 7), demonstrating therefore that miR-199a-5p promotes TGF-β signaling through Cav1 regulation (Fig. 4, E to H and fig. S7D). Overall, these results provide important *in vivo* mechanistic insights into the profibrotic role of miR-199a-5p during lung fibrogenesis, and strongly suggest that targeting this miRNA may be a valid therapeutic option in IPF.

**Fig. 4.**
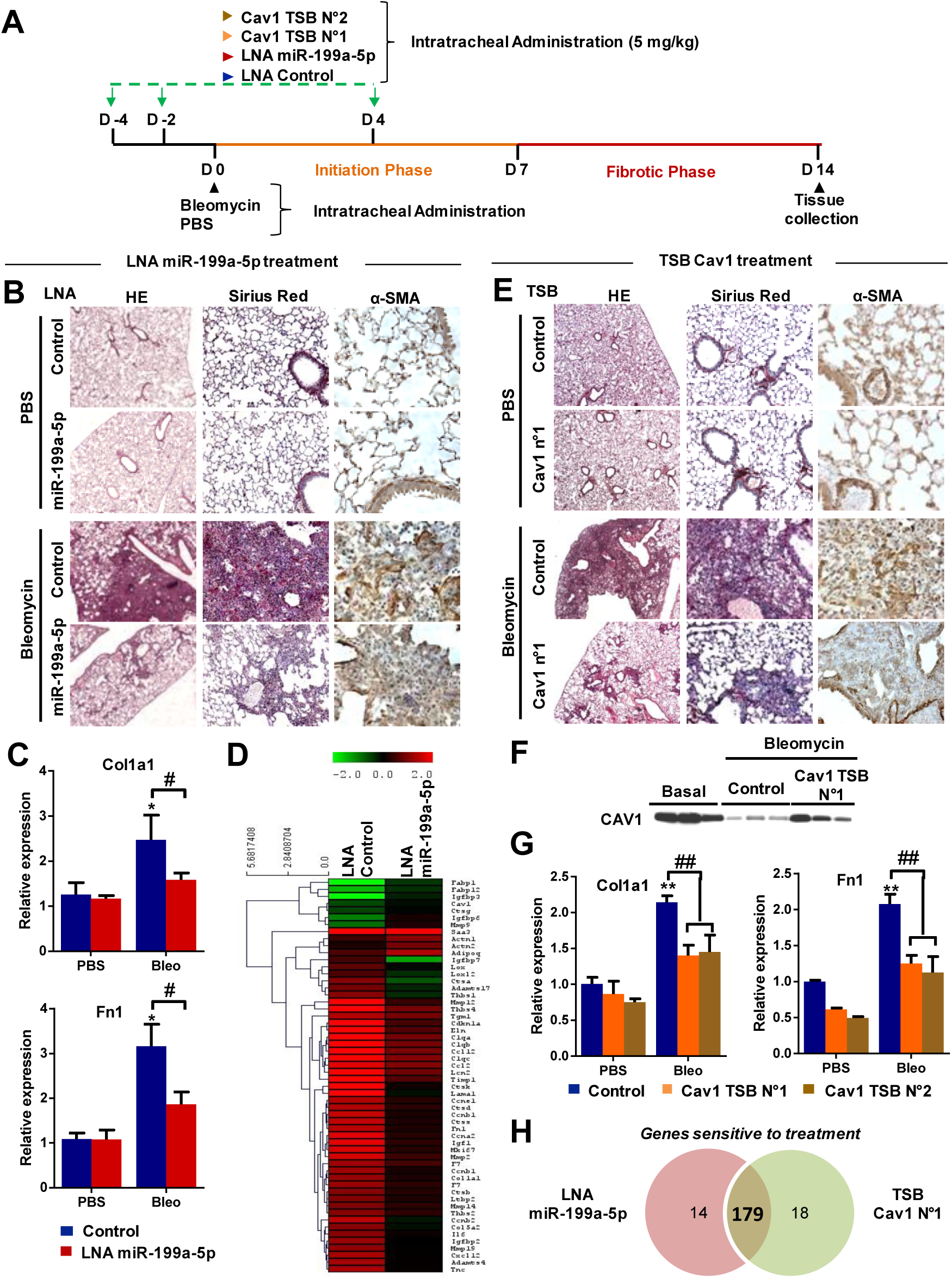
CAV1 represents the main target associated with miR-199a-5p profibrotic activity in bleomycin-induced lung fibrosis mouse model. **(A)** Diagram showing the experimental protocol used. Mice (*n*=4–6 in each group) received control LNA-probes, Cav1 TSBs or miR-199a-5p antisense LNA-probes intratracheally 2 days and 4 days before intratracheal instillation of bleomycin or PBS as well as 4 days following bleomycin treatment. 14 days after bleomycin administration, mice were sacrificed and lungs were collected for further analysis **(B,D)** Silencing of miR-199a-5p prevents bleomycin-induced lung fibrosis in mice. **(B)** Paraffin sections were prepared from lungs of mice that received LNA-miR-199a-5p inhibitor or control probe and immunohistochemical analysis of α-SMA expression was performed as well as H&E and sirius red staining at x25, x10 and x2.5 magnification respectively. One representative section is shown for each condition. **(C)** Bar charts showing the relative pulmonary expression of Col1a1 and Fn1 (compared to Ppia) in the LNA-miR-199a-5p protocol. Data are expressed as mean ± SEM. * *P*<0.05, # *P*<0.05. **(D)** Heatmap showing the effect of LNA-miR-199a-5p and LNA-control treatment on bleomycin pulmonary response, expressed as log2 ratio (Bleo / PBS) on a subset of genes associated with lung fibrosis (Dataset 6). **(E-H)** miR-199a-5p profibrotic activity relies on Cav1 targeting *in vivo*. **(E)** Paraffin sections were prepared from lungs of mice that received Cav1 TSB or control probe and immunohistochemical analysis of α-SMA expression was performed as well as H&E and sirius red staining at x25, x10 and x2.5 magnification respectively. One representative section is shown for each condition. **(F)** Western Blot showing that specifically inhibiting miR-199a-5p binding on mouse Cav1 3’UTR using Cav1 TSB inhibits bleomycin-induced downregulation of CAV1 in mouse lungs. **(G)** Bar charts showing the relative expression of Col1a1 and Fn1 in lungs from mice that received the two distinct Cav1 TSBs or control probe, compared to Ppia. Data are expressed as mean ± SEM. ** *P*<0.01, ## *P*<0.01. **(H)** Venn diagram showing an overlap between genes significantly modulated in response to bleomycin in the LNA-miR-199a-5p (Dataset 6) and the Cav1 TSB protocol (Dataset 7). The number of bleomycin-modulated transcripts affected by the 2 molecules *in vivo* are shown.

### Systemic administration of LNA-based inhibitor of miR-199a-5p diminishes the severity of bleomycin-induced lung fibrosis in mice without causing adverse effects

To explore whether inhibition of miR-199a-5p is a potential therapeutic option for the treatment of lung fibrosis, we repetitively administered miR-199a-5p antisense probes i.p. from day 8 after intratracheal instillation of bleomycin, a time when inflammatory responses start to subside and active fibrogenesis occurs *(22, 33)*, and then analyzed the extent of lung fibrosis 18 days following bleomycin treatment (Fig. 5A). Under these conditions, LNA-mediated silencing of miR-199a-5p attenuated lung collagen deposition as assessed by hydroxyprolin quantification (Fig. 5B). Finally, the therapeutic potential of miR-199a-5p silencing was further defined by evaluating its safety profile. Importantly, systemic administration of miR-199a-5p inhibitor in mice did neither induce acute nor chronic toxicity especially in liver and kidneys (fig. S8). Altogether, these results provide preclinical proof-of-concept for targeting miR-199a-5p as a new effective and safe approach to treat lethal pulmonary fibrosis.

**Fig. 5.**
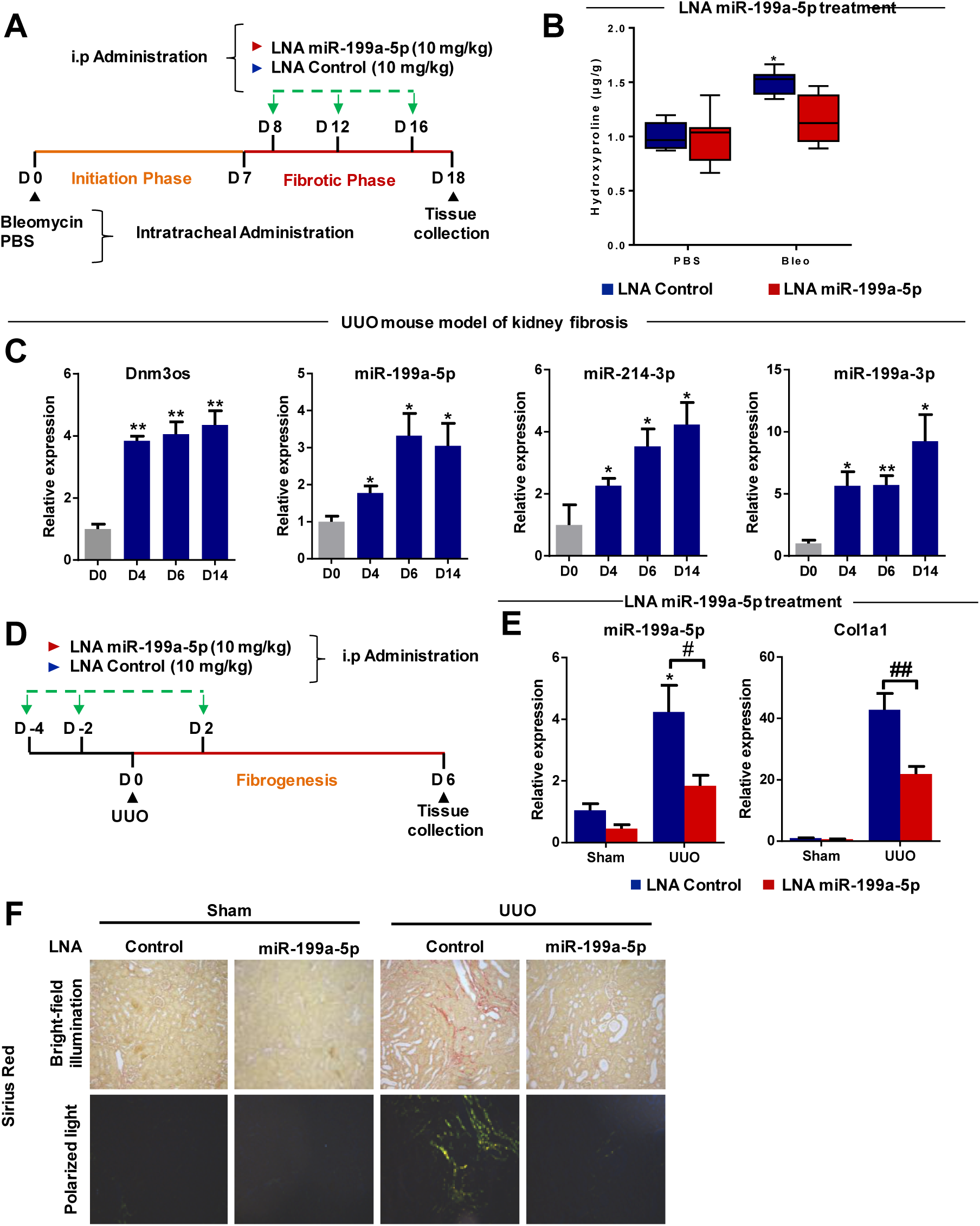
Systemic administration of LNA-miR-199a-5p inhibitor in mice diminishes the severity of bleomycin-induced lung fibrosis and ameliorates kidney fibrosis. **(A)** Diagram showing the experimental protocol used to study lung fibrosis. Mice (*n* = 6–10 in each group) were given bleomycin intratracheally. On day 8, 12, and 16 after bleomycin administration, control or LNA-199a-5p antisense probes were injected intraperitoneally. 18 days after bleomycin instillation, mouse lungs were collected. **(B)** Box plots showing whole lung collagen content measured by hydroxyproline quantification. The box represents the 25–75% quartiles, the line in the box represents the median and whiskers represent the range. * *P*<0.05. **(C-F)** Impact of miR-199a-5p silencing on kidney fibrosis. **(C)** Bar charts showing the relative expression of Dnm3os and its associated miRNAs during kidney fibrogenesis. (*n*=3-5 mice per group) * *P*<0.05. **(D)** Diagram showing the experimental protocol used to study renal fibrosis. Mice (*n*= 3-5 mice in each group) received control LNA-probes or miR-199a-5p antisense LNA-probes intraperitoneally 2 days and 4 days before unilateral ureteral obstruction (UUO) as well as 2 days following surgery. 6 days after UUO, mice were sacrificed and kidneys were collected for further analysis. **(E)** Bar charts showing the relative renal expression of miR-199a-5p (compared to SNO251) as well as Col1a1 (compared to Ppia) according to the protocol described above. Data are expressed as mean ± SEM. * *P*<0.05, # *P*<0.05, ## *P*<0.01. **(F)** Paraffin sections were prepared from kidneys of mice that received LNA-199a-5p or control probe and stained with sirius red. One representative section viewed under bright field or polarization contrast is shown for each condition at x10 magnification.

### Inhibition of miR-199a-5p ameliorates kidney fibrosis

Finally, given the consistent upregulation of miR-199a-5p that has been reported in various fibroproliferative diseases *(21, 34, 35)*, we also investigated whether targeting this miRNA may benefit in fibrotic conditions other than those affecting the lung. For this, we used the Unilateral Ureteral Obstruction (UUO) mouse model, which induces a progressive renal fibrosis, similar to that seen in most human progressive renal diseases *(36)*. As expected, expression of Dnm3os and its associated miRNAs increased during kidney fibrogenesis as early as 4 days following kidney injury (Fig. 5C). Consistent with our *in vivo* findings obtained in lung fibrosis, treatment with LNA–modified antisense probes designed against miR-199a-5p (Fig. 5D) prevented the enhanced renal expression of miR-199a-5p and efficiently attenuated Col1a1 gene expression 6 days after UUO (Fig. 5E). Moreover, sirius red staining of kidney sections demonstrated a striking attenuation of collagen deposition in kidney of mice pretreated with LNA-199a-5p (Fig. 5F). Altogether, these results strongly suggest that miR-199a-5p represents a potential therapeutic target in various refractory fibroproliferative disorders.

## Discussion

The traditional view of genome organization has radically changed in the last decade with the discovery of vast amounts of functional RNA transcripts with no protein-coding potential *(37)*. As mechanistic studies have linked these ncRNAs to a broad spectrum of complex human disorders, ncRNA-based therapies offer new tremendous opportunities to treat incurable diseases *(18, 32)*. In line with this, we reasoned that ncRNAs play a substantial role in the pathogenic events leading to fibrogenesis and therefore may represent new valuable druggable targets especially for the treatment of lethal fibrotic diseases. In this study, we focused on IPF, a devastating fibroproliferative lung disorder, for which solid preclinical and clinical evidence support antagonism of the TGF-β pathway as a powerful therapeutic modality in particular to reduce (myo)fibroblast activity and consequently pulmonary fibrogenesis *(1, 8)*. Accordingly, the identification of profibrotic ncRNAs mediating TGF-β-induced lung fibroblast activation may have strong therapeutic implication, especially as TGF-β-targeted therapies have not yet reached the clinic *(8, 9)*. Our results showed that TGF-β profoundly influences the ncRNA transcriptional program of lung fibroblasts, suggesting that non-coding transcripts are likely to exert important regulatory functions of TGF-β signaling rather than being passive by-products of transcriptional noise. In particular, we identified the lncRNA DNM3OS, as a critical downstream effector of the TGF-β Pathway in lung fibroblasts.

DNM3OS is an antisense transcript, located within an intron of the human dynamin-3 (DNM3) gene. As this RNA contains only short and poorly conserved open reading frames, it is classified as a long non-coding RNA by virtue of being more than 200 nucleotides in length *(15)*. In contrast to most lncRNAs, DNM3OS biological function has been experimentally defined and involves post-transcriptional regulation of gene expression *(20, 38)*. Indeed, DNM3OS locus contains two highly conserved miRNA genes, miR-199a-2 and miR-214, which when processed into their mature forms, repress expression of their target genes to induce biological effects. Consistent with this, our sequencing data showed that expression of DNM3OS in lung fibroblasts stimulated with TGF-β mirrors that of the mature miRNAs encoded by the miR-199~214 cluster. While miR-199a-5p is an established critical effector of TGF-β signaling in lung fibroblasts *(21)*, whether the other clustered miRNAs also contribute to TGF-β profibrotic activity is unclear. Therefore, we hypothesized that DNM3OS is a reservoir of fibromiRs that could be rapidly mobilized following TGF-β stimulation, to synergistically promote lung fibroblast activation by targeting distinct components of the TGF-β pathway (Fig. 6). Indeed, given their co-expression, clustered miRNAs are known to jointly regulate molecular pathway either by co-targeting individual genes or by targeting different components of the same pathway *(16)*. In line with this, our results demonstrated that processing of DNM3OS transcript in lung fibroblasts give rise to 3 individual profibrotic miRNAs, miR-199a-5p/3p and miR-214-3p, which function as critical intermediate of TGF-β signaling by targeting distinct functionally related genes. Mechanistically, we showed that these fibromiRs influence TGF-β signaling in a multifaceted way, through two distinct mode of action consisting of either signal amplification or mediation by respectively establishing positive feed forward loop or acting as essential downstream signal effectors. Indeed, TGF-β-induced upregulation of miR-199a-5p results in the inhibition of a negative feedback mechanism involving CAV1, leading to both SMAD and non-SMAD signaling amplification. In contrast, our results ascribed to miR-199a-3p a specific role in mediating TGF-β-induced suppression of HGF/KGF secretion, two growth factors potently promoting tissue repair *(29, 30)*. Lastly, we showed that miR-214-3p is specifically involved in the mediation and promotion of the GSK3β/β-catenin pathway, a non-SMAD component of the TGF-β signaling cascade, by respectively targeting GSK-3β and COX-2. Collectively, these results unambiguously establish DNM3OS as a *bona fide* profibrotic lncRNA, which specifically regulates multiple fibrogenic components of the TGF-β pathway in lung fibroblasts, by serving as a precursor of fibromiRs.

**Fig. 6.**
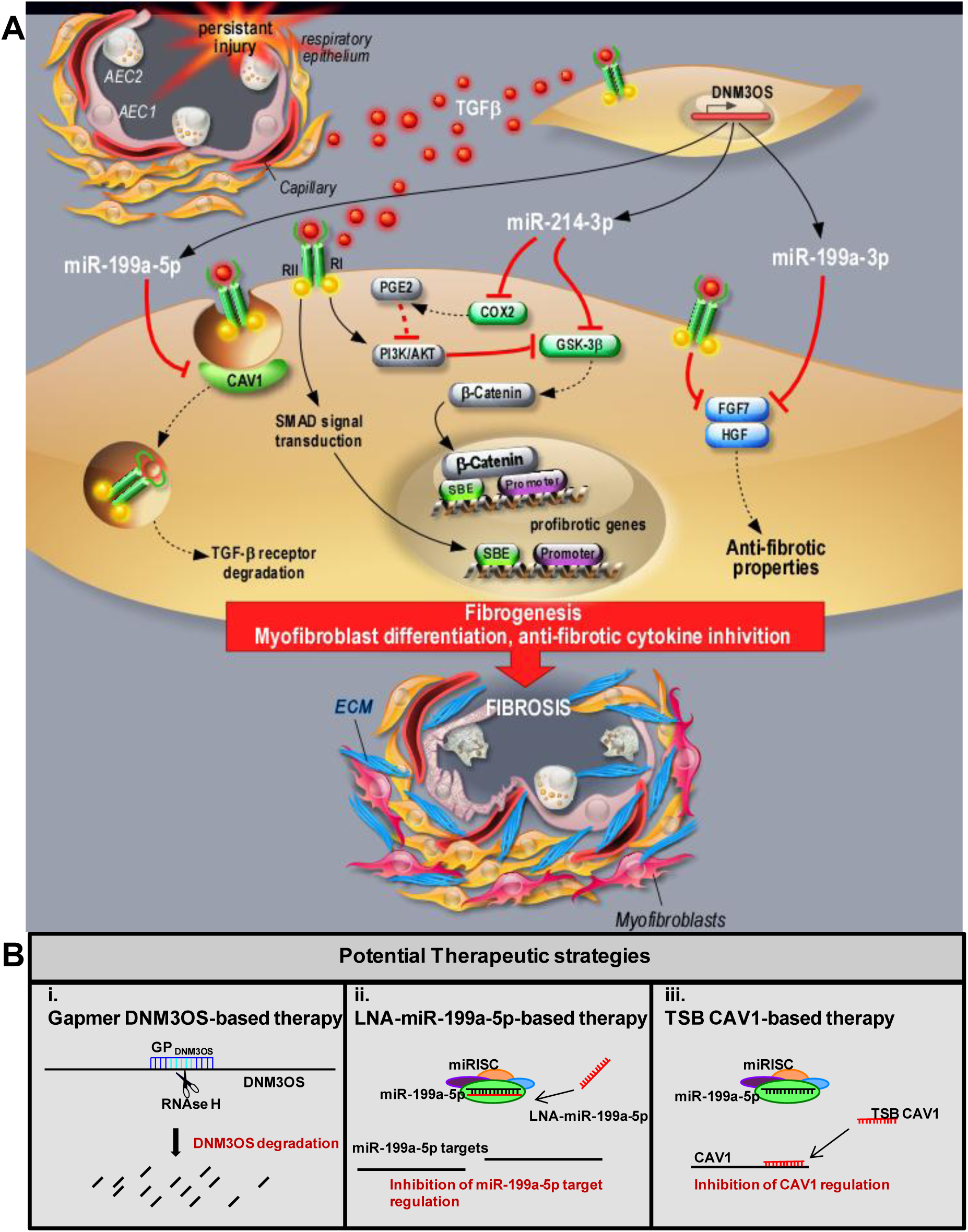
Proposed model of DNM3OS profibrotic function in lung fibrogenesis and potential strategies for therapy. **(A)** Proposed model. Persistent injury of the respiratory epithelium causes release of profibrotic factors such as TGF-β. In lung fibroblasts, TGF-β binds to TGF-βR I (RI) and II (RII) and increases expression of DNM3OS and its associated mature miRNAs, miR-199a-5p, miR-199a-3p and miR-214-3p. Production of miR-199a-5p in response to TGF-β in lung fibroblasts results in CAV1 downregulation and subsequently, impaired TGF-β/TGF-βR complex degradation. This miRNA-mediated mechanism for low CAV1 expression promotes both canonical and non-canonical TGF-β signaling pathways and the pathogenic activation of lung fibroblasts. The increased expression of miR-214-3p promotes the non-canonical GSK-3β/β-catenin axis of TGF-β signaling by targeting COX-2 and GSK-3β resulting in myofibroblast differentiation. The upregulation of miR-199a-3p mediates TGF-β-induced inhibition of both FGF7 and HGF release. AEC: Alveolar Epithelial Cell; TGF-β: Transforming Growth Factor β; DNM3OS: Dynamin 3 Opposite Strand; CAV1: Caveolin 1; COX-2: Cyclooxygenase 2; ECM: Extracellular Matrix; PGE2: Prostaglandin E 2; PI3K: Phosphatidylinositol 3 Kinase; GSK-3β: Glycogen Synthase Kinase 3 β; SBE: SMAD Binding Element; FGF7: Fibroblast Growth Factor 7; HGF: Hepatocyte Growth Factor. **(B)** Potential therapeutic strategies to repress TGF-β profibrotic signaling: (i) RNase H-mediated silencing of DNM3OS, (ii) inhibition of miR-199a-5p (iii) preventing miR-199a-5p binding to CAV1.

Although notable progress has been made in the comprehension of TGF-β signaling, the development of anti-fibrotic drug targeting this growth factor has lagged behind, largely due to our poor understanding of the precise molecular mediators driving TGF-β profibrotic effects *(8, 9)*. In this study, we provide new insights into the molecular circuitry underlying TGF-β-induced profibrotic signaling in lung fibroblasts, and identify the lncRNA DNM3OS as a new attractive target for anti-fibrotic drug development. Currently, the most efficient approach to therapeutically disrupt nuclear lncRNA function relies on RNase H-mediated silencing of target RNA using chemically modified antisense oligonucleotides such as gapmer *(19)*. Using a gapmer targeting DNM3OS, we could indeed efficiently inhibit both DNM3OS transcript expression and the production of the 3 mature miRNAs *in vitro* in human cells. This represents to our knowledge the first proof-of-concept for the use of gapmers to target a nuclear miRNA precursor and silence a whole cluster of miRNAs. Nevertheless, the therapeutic potential of DNM3OS targeting *via* RNAse H raises several conceptual and technical issues. First, gapmer design algorithms mostly retrieved human target sequences that are not evolutionary conserved, making pre-clinical testing of these molecules challenging. Second, as Dnm3os has been shown to be indispensable for normal growth *(20)*, *in vivo* silencing of this non-coding transcript may cause significant toxicity. Last, pharmacological modulation of lncRNA, in contrast to miRNA, is still in its infancy and requires further preclinical evaluation before entering clinical trials. Therefore, we reasoned that individual targeting of profibrotic miRNAs processed from DNM3OS would be a more suitable alternative from a translational point of view, given their high degree of sequence conservation and the recent clinical success of miRNA-based therapy *(39)*. In light of our *in vitro* findings, we chose to focus on miR-199a-5p, whose overexpression in lung fibroblasts most faithfully recapitulates the complexity of TGF-β pathway by affecting both SMAD and non-SMAD signaling. Our *in vivo* data not only confirms the deleterious role of miR-199a-5p in lung fibrogenesis, but also demonstrates that this miRNA-mediated pro-fibrotic effect is largely explained by a single target interaction with CAV1. Nevertheless, while selectively interfering with CAV1 targeting warrants consideration in lung fibrosis, especially to avoid potential unintended effects resulting from miR-199a-5p silencing, this may not be true in other organ systems. Indeed, although CAV1 anti-fibrotic role has been widely described in lung *(21, 40, 41)*, whether it also has a beneficial role in other fibrotic conditions is less clear, especially in liver *(42)*. In addition, the interaction between CAV1 and miR-199a-5p is not established in fibrotic diseases other than pulmonary fibrosis. In contrast, miR-199a-5p is consistently upregulated in fibrotic diseases regardless of the injured organ *(21, 34, 35)*, and thus represents a general anti-fibrotic target. Therefore, we focused our attention on miR-199a-5p silencing and further showed that LNA-mediated silencing of this miRNA not only prevents lung fibrosis development but also ameliorates kidney fibrosis and improves established pulmonary fibrosis. Importantly, we also demonstrated the apparent safety of antimiR-199a-5p systemic treatment, as the clinical chemistry and pathologic analyses demonstrated no evidence of toxicity, in particular in kidney and liver. Indeed, given their predominant accumulation in liver and kidney, one major concern of antimiR therapy is to achieve sufficient intracellular concentrations of oligonucleotides in the target tissue to evoke a therapeutic effect without causing liver and renal toxicity *(43)*.

In conclusion, the results of this study further highlight the pivotal roles played by ncRNAs in mediating changes in gene expression and cell functions occurring during pulmonary fibrosis. In particular, our results identified DNM3OS as a new determinant of pulmonary fibrosis and mechanistically ascribed its profibrotic effect to the regulation of the complex molecular events leading to TGF-β-dependent activation of lung fibroblasts, in particular by serving as a precursor of 3 distinct fibromiRs. We thus anticipate that strategies surrounding DNM3OS targeting, especially miR-199a-5p silencing, may represent a new effective therapeutic option to treat IPF and possibly other lethal fibrotic diseases.

## Materials and Methods

### General experimental approaches

Sample size was chosen empirically based on our previous experiences in the calculation of experimental variability; no statistical method was used to predetermine sample size and no samples, mice or data points were excluded from the reported analyses. Experiments were performed non-blinded using treatment group randomized mice.

### Animal treatment

All animal care and experimental protocols were conducted according to European, national and institutional regulations (Protocol Numbers: CEEA 142012 and CEEA 162011: University of Lille; 00236.03: CNRS). Personnel from the laboratory carried out all experimental protocols under strict guidelines to insure careful and consistent handling of the mice. All *in vivo* experiments were performed using 9–12 weeks old male C57BL/6 mice purchased from Charles River.

### Mouse model of lung fibrosis

To induce fibrotic changes, 50 μL bleomycin (1Unit/kg) or PBS were aerosolized in mouse lungs using a MicroSprayer Aerosolizer (Penn-Century Inc). LNA-modified oligonucleotides used for *in vivo* experiments were purchased from Exiqon, dissolved in PBS and then injected intratracheally using a MicroSprayer Aerosolizer (5 mg/kg) or intraperitoneally using an insulin syringe (probes are listed in table S6).

### Mouse model of kidney fibrosis

Mice underwent anesthesia by intraperitoneal injection of pentobarbital (50 mg/kg). After a standard laparotomy, the left proximal ureter was exposed and ligatured with 4-0 silk at two points. The sham operation consisted of a similar identification of the left ureter, but ligature of the ureter was not performed.

### Histopathology

Mouse lungs, livers and kidneys were fixed overnight with neutral buffered formalin and then embedded in paraffin. 5-micrometer-thick sections were mounted and stained with hematoxylin and eosin as well as Sirius Red to assess the degree of fibrosis.

### Mouse blood chemistry

Glucose, blood urea nitrogen (BUN), alkaline phosphatase (ALP), alanine aminotransferase (ALAT), aspartate aminotransferase (ASAT), cholesterol, lipase, triglycerides, and creatine phosphokinase (CPK) were measured on plasma samples using a COBAS 8000 fully automated analyzer.

### Hydroxyproline quantification

Lung hydroxyproline content was assayed as previously described*(44)*. Briefly, the right lung of each mouse was dissected free of extraneous tissue, homogenized in 2 mL of water, and then incubated in 0.5% trichloroacetic acid on ice for 20 min. After centrifugation, the pellet was resuspended once in 1.0 mL of 12 N HCl and baked at 110°C for 24 hours, and then in 2.0 mL of distilled water. Fifty microliters of each sample were incubated for 20 min at room temperature in 100 mL of a solution containing 1.4% chloramine-T, 0.5M sodium acetate, and 10% isopropanol, before addition of 100 mL of Ehrlich’s solution (1.0M p-dimethylaminobenzaldehyde dissolved into 70% isopropanol plus 30% perchloric acid). Optical density was measured after 15 min incubation at 65°C. Purified hydroxyproline standards were used to convert optical density values to hydroxyproline concentrations.

### Immunohistochemistry

5-µm paraffin-embedded sections were sequentially incubated in xylene (5 min twice), 100% ethyl alcohol (5 min twice), 95% ethyl alcohol (5 min twice), and 80% ethyl alcohol (5 min). After washing with water, the sections were antigen-retrieved using citrate buffer (pH 6.0; DAKO) in a domestic microwave oven for 20 minutes and cooled to ambient temperature. Sections were then washed with TBS-T and quenched with 3% hydrogen peroxide in TBS for 10 min, blocked for avidin/biotin activity, blocked with serum-free blocking reagent, and incubated with primary antibody (α-SMA, 1:500). Immunohistochemical staining was developed using the DAB substrate system (DAKO).

### Cell lines, reagents, and antibodies

Normal human pulmonary fibroblasts MRC-5 (CCL-171), and HEK-293 (CRL-1573) cells were purchased from the American Type Culture Collection (ATCC), frozen at an early passage. Each vial used for experiments was cultured for a limited number of passages (<15). For maintenance, cells were cultured in the appropriate medium (MEM for MRC-5, DMEM for HEK-293) containing 10% fetal calf serum (FCS), 1% Penicillin Streptomycin, 1% glutamax, at 37°C with 5% v/v CO2. Regular testing for mycoplasma contamination was carried out by PCR. Recombinant TGF-β and TNF-α were purchased from Peprotech and used at 10 ng/mL unless indicated. Recombinant Wnt 3A was purchased from R&D Systems and used at 150 ng/mL. PGE2 was purchased from Tocris Bioscience and used at 10nM. The following antibodies were used: goat anti-HSP60 (sc-1052, Santa Cruz Biotechnology Inc.), rabbit anti-lamin (ab106682, abcam), rabbit anti-CAV1 (sc-894, Santa Cruz Biotechnology Inc.), rabbit anti-β-tubulin (2146), COX-2 (12282), SMAD4 (9515), SMAD2 (5339), p-SMAD2 (4060), Akt (9272), p-Akt (4060) GSK-3β (12456), p-GSK-3β (5558) and anti-β-Actin (13E5) (Cell Signaling), mouse anti-αSMA (1A4, Dako) for immunohistochemistry and (4A8-2H3, Abnova) for Western Blot and immunocytofluorescence, anti-β-Catenin mAb (610153, BD Biosciences).

### Transfection and luciferase assays

Pre-miRNAs and control miRNA (miR-Neg # 1) were purchased from Ambion. Target site blockers, gapmers, LNA-based miRNA inhibitors and their respective controls were ordered from Exiqon. siRNA directed against GSK3β and control siRNA (Silencer Select validated siRNAs) were purchased from Thermo Fisher Scientific. MRC5 cells were grown in 10% FCS in DMEM and transfected at 30 to 40% confluency in 6-24 well plates using Lipofectamin RNAi MAX™ (Thermo Fisher Scientific) with pre-miRNAs, siRNAs, LNA inhibitors and target site blockers at a final concentration of 10nM unless indicated. Molecular constructs were made in psiCHECK-2 (Promega) by cloning annealed oligonucleotides derived from 3′ UTR target genes (table 7) upstream of the Renilla luciferase gene using the XhoI and NotI restriction sites. HEK293 cells were plated into 96-well plates and cotransfected using lipofectamin 2000 (Thermo Fisher Scientific) with 0.2 µg of psiCHECK-2 plasmid constructs and pre-miRNAs or control pre-miRNA. 48 hours after transfection, Firefly and Renilla luciferase activities were measured using the Dual-Glo Luciferase assay (Promega).

### Protein extraction and immunoblotting

Cells or tissues were lysed in RIPA lysis buffer containing protease and phosphatase inhibitors cocktail (Pierce). Nuclear and cytoplasmic fractionations were performed using NE-PER Nuclear and Cytoplasmic Extraction kit according to manufacturer’s instructions (Pierce). Lysates were quantified for protein concentrations using the BCA Protein Assay Kit (Pierce). Proteins were separated by SDS-polyacrylamide gel and transferred onto PVDF membranes (GE Healthcare). The membranes were blocked with 5% fat free milk or BSA (in case of phospho protein) in Tris-buffered saline (TBS) containing 0.1% Tween-20 (TBS-T) and subsequently incubated with their respective primary antibodies overnight at 4°C (all antibodies were used at 1:1000 except HSP60 which was used at 1:4000). After washing with TBS-T for 30 minutes at room temperature, the membrane was further incubated with horseradish peroxidase–conjugated secondary antibodies for 1.5 hours, followed by 30 minutes of washing with TBS-T. Protein bands were visualized with Amersham ECL select substrates (GE Healthcare). The original blots are shown in fig. S8 to S10.

### Assessment of KGF, HGF and PGE2 release from lung fibroblasts

MRC5 cells were plated into 6-well plates and transfected as described above with pre-miRNAs, LNA-based miRNA inhibitors or negative control. Twenty-four hours after transfection, cells were stimulated with TGF-β or TNF-β. 24h after stimulation, supernatants were collected and KGF, HGF and PGE2 were quantified by ELISA kits purchased from R&D Systems according to manufacturer’s protocol.

### Immunocytofluorescence analysis

MRC5 cells were grown on a Round Glass Coverslips Ø 16 mm (Thermo Fisher Scientific) placed inside a 12-well Plate. Coverslips were washed in phosphate-buffered saline and fixed in 4% paraformaldehyde for 15 min, cells were then permeated using 0.1% Triton X-100 (Agilent Technologies) for 10 min and blocked with PBS solution containing BSA (3%) for 30 min. Incubation with primary antibodies was performed in a blocking solution BSA (1%) at 37°C for 1 h at the following dilutions: α-SMA (1:1000), β-catenin (1:500). After three washes with PBS, cells were incubated with secondary Alexa Fluor 488 goat anti-Mouse IgG (Invitrogen) (1:500), Alexa Fluor 647 goat anti-rabbit IgG (Thermo Fisher Scientific) (1:500) and Alexa Fluor 647 Phalloidin (A22287 - Life technologies) (1Unit/slide). Forty-five min later, coverslips were mounted on microscope slides using ProLong Gold Antifade Reagent with DAPI (Thermo Fisher Scientific). Fluorescence was viewed using a FV10i Olympus confocal scanning microscope.

### Invasion assay

Invasion of MRC5 fibroblasts was assessed using commercially available 24-well BioCoat Matrigel Invasion Chamber (BD Biosciences). In brief, pulmonary fibroblasts were transfected either with pre-miRNAs, gapmer, LNA-based miRNA inhibitors or negative controls as described above. Twenty-four hours after transfection, cells were harvested with trypsin-EDTA, centrifuged, and resuspended in MEM medium. Cell suspensions (1×10^5^ cells/well) were added to the upper chamber. Bottom wells of the chamber were filled with DMEM medium containing 10% FCS as chemoattractant, whereas the upper chamber was filled with either MEM or MEM plus TGF-β (10ng/mL). After incubation for 48 h at 37°C, the non-invading cells on the top of the membrane were removed with a cotton swab. Membrane containing invading-cells were fixed with methanol, washed three times with PBS and mounted with DAPI hard set (Vector Laboratories) onto glass slides for fluorescent microscopy.

### RNA isolation

Total RNAs were extracted from tissues and cell samples using TRIzol reagent (Invitrogen). Integrity of total RNAs was assessed using an Agilent BioAnalyser 2100 (Agilent Technologies) (RIN above 7).

### Gene expression profiling

#### RNA-seq

Libraries were generated from 500ng of total RNAs using Truseq Stranded Total RNA kit (Illumina). Libraries were then quantified with KAPA library quantification kit (Kapa biosystems) and pooled. 4nM of this pool were loaded on a high output flowcell and sequenced on a Illumina NextSeq500 sequencer using 2×75bp paired-end chemistry for a total of 378M reads. Reads were aligned to the human genome release hg19 with STAR 2.5.2a with following parameters “--outFilterIntronMotifs RemoveNoncanonicalUnannotated --alignMatesGapMax 1000000 --outReadsUnmapped Fastx --alignIntronMin 20 --alignIntronMax 1000000 --alignSJoverhangMin 8 --alignSJDBoverhangMin 1 --outFilterMultimapNmax 20”. Quantification of genes was then performed using featureCounts release subread-1.5.0-p3-Linux-x86_64 with “--primary -g gene_name -p -s 1 -M -C” options based on Ensembl GTF release 75 annotations.

#### Small RNA-Seq

500ng of total RNA were ligated, reverse transcribed and amplified (18 cycles) with the reagents from the NextFlex small RNAseq kit V3 (Bioo scientific). Amplified libraries were quantified with the Bioanalyzer High Sensitivity DNA Kit (Agilent), pooled and size-selected from 140 nt to 170 nt with the LabChip XT DNA 300 Assay Kit (Caliper Lifesciences). Libraries were then sequenced on an Illumina Nextseq 500 Mid Flowcell with 75pb reads for a total of 147M reads.

*Analysis*: Cutadapt (v1.2.1) was used to remove Illumina 3p adaptor with parameter “--adapter=TGGAATTCTCGGGTGCCAAGG”. Trimmed reads under 15 bases were discarded from further analysis. Mapping was done with Bowtie2 versus human hg19 Ensembl release GRCh37.75 builds using “--local --very-sensitive-local -k 24” parameters. Count table was generated based on GTF annotations files from mirbase database (v21), Ensembl ncrna databases (rel73), tRNAs for tRFs counting coming from UCSC table browser and piRNA clusters from piRNAclusterDB..

#### Expression microarrays

For gene expression arrays, RNA samples were labeled with Cy3 dye using the low RNA input QuickAmp kit (Agilent) as recommended by the supplier. 825 ng of labeled cRNA probes were hybridized on 8×60K high density SurePrint G3 gene expression mouse or human Agilent microarrays.

#### Statistical analysis and Biological Theme Analysis

Microarray data analyses were performed using R (http://www.r-project.org/). Quality control of expression arrays was performed using the Bioconductor package arrayQualityMetrics and custom R scripts. Additional analyses of expression arrays were performed using the Bioconductor package limma. Briefly, data were normalized using the quantile method. No background subtraction was performed. Replicated probes were averaged after normalization and control probes removed. Statistical significance was assessed using the limma moderated t-statistic Quality control of RNA-seq and Small RNA-seq count data was assessed using in-house R scripts. Normalization and statistical analysis were performed using Bioconductor package DESeq2. All P-values were adjusted for multiple testing using the Benjamini-Hochberg procedure, which controls the false discovery rate (FDR). Differentially expressed genes were selected based on an adjusted p-value below 0.05. Enrichment in biological themes (Molecular function, Upstream regulators and canonical pathways) and biological networks analysis were performed using Ingenuity Pathway Analysis software (http://www.ingenuity.com/). Gene Set Enrichment Analysis (GSEA)*(25)* was used to determine whether an *a-priori* defined set of genes can characterize differences between two biological states. Hierarchical clusterings were done with the MultiExperiment Viewer (MeV) program version 4.9, using a Manhattan distance metric and average linkage.

#### MiRNA targets analysis

MiRonTop is an online java web tool (available at http://www.genomique.info/) that integrates whole transcriptome expression data (i.e. microarray or sequencing) to identify the potential implication of miRNAs on a specific biological system. Briefly, MiRonTop ranks the transcripts into 2 categories (‘Upregulated’ and ‘Downregulated’), according to thresholds for expression level, differential expression and statistical significance*(45)*. It then calculates the number of predicted targets for each miRNA, according to the prediction software selected (Targetscan, exact seed search, TarBase), in each set of genes. Enrichment in miRNA targets in each category is then tested using the hypergeometric function.

#### Microarray Datasets from IPF lung-derived samples

Publicly available microarray data from patients with IPF was obtained from the Gene Expression Omnibus (GEO, http://www.ncbi.nlm.nih.gov/geo/). We used the gene set IDs GSE32537 and GSE21394 to calculate the differential pulmonary expression of DNM3OS and its associated mature miRNAs respectively.

### RNA FISH

Custom Stellaris^®^ FISH Probes were designed against DNM3OS using the Stellaris^®^ RNA FISH Probe Designer (Biosearch Technologies) available online at www.biosearchtech.com/stellarisdesigner (version 4.2). MRC5 cell were grown on µ-Dish 35mm glass bottom (Ibidi), and stimulated with TGF-β at a final concentration of 10 ng/mL. 24h after exposure, fixed cells were permeabilized using 70% ethanol and were hybridized with the DNM3OS Stellaris RNA FISH Probe set labeled with CAL Fluor Red 590 (Biosearch Technologies), following the manufacturer’s instructions at the final concentration of 500 nM available online at www.biosearchtech.com/stellarisprotocols. Acquisition was performed using an inverted confocal microscope LSM 710 (Zeiss) with the high-resolution module AiryScan (Zeiss) and a 63x/1.4 oil immersion lens. Images were processed using Zen software (Zeiss).

### Quantitative RT–PCR

#### Mature miRNA expression

MiRNA expression was assessed using TaqMan MicroRNA Reverse Transcription Kit and TaqMan MicroRNA Assays (Thermo Fisher Scientific) as specified by the manufacturer. Real-time PCR was performed using Universal Master Mix II (Thermo Fisher Scientific) and ABI 7900HT real-time PCR machine. Expression levels of mature microRNAs were evaluated using comparative CT method. For normalization, transcript levels of RNU44 (human samples) and SNO251 (mouse samples) were used as endogenous control for miRNA real time PCR. Assays are listed in table 8.

#### Gene expression

Reverse transcription was performed using High-Capacity cDNA Reverse Transcription Kit (Thermo Fisher Scientific). Expression levels of both human and mouse genes (primers listed in table S9) were performed using Fast SYBR Green Master Mix (Thermo Fisher Scientific) and ABI 7900HT real-time PCR machine. Expression levels were evaluated using comparative CT method. For normalization, transcript levels of PPIA (human and mouse samples) were used as endogenous control for gene expression.

### Statistical analysis

All data were graphed using GraphPad Prism. Results are given as mean±S.E.M. Statistical analyses were performed using the two-tailed Mann–Whitney test for single comparison; the variance was similar in the groups being compared. In each case, a *P* value <0.05 was considered statistically significant.

### Data-availability

Expression datasets that support the findings of this study have been deposited in the Gene Expression Omnibus SuperSerie record GSE97834 containing 7 distinct datasets under the following accession codes:

- Dataset 1: GSE97829. RNA-Seq analysis of human lung fibroblasts exposed to TGF-β.
- Dataset 2: GSE97832. Small RNA-Seq analysis of human lung fibroblasts exposed to TGF-β.
- Dataset 3: GSE97823. Impact of DNM3OS silencing on human lung fibroblast response to TGF-β.
- Dataset 4: GSE97824. Impact of miR-199-5p, miR-199a-3p and miR-214-3p 654 overexpression on human lung fibroblasts.
- Dataset 5: GSE97833. TGF-β signature of MRC-5 fibroblasts.
- Dataset 6: GSE97825. Impact of miR-199a-5p silencing on bleomycin-induced lung fibrosis.
- Dataset 7: GSE97826. Impact of CAV1 target site blocker (CAV1 TSB) on bleomycin-induced lung fibrosis.

## Supplementary Materials

Fig. S1. The non-coding transcriptome is a key component of the TGF-β response of lung fibroblasts

Fig. S2. Identification of DNM3OS as a key effector of TGF-β signaling.

Fig. S3. DNM3OS is processed into three profibrotic miRNAs in lung fibroblasts.

Fig. S4. Identification of the cellular pathways and gene targets associated with the miR-199a~214 cluster in human lung fibroblasts.

Fig. S5. miR-214-3p mediates TGF-β-induced lung fibroblast activation by targeting two distinct targets: GSK-3β and COX-2.

Fig. S6. KGF/FGF7 and HGF are direct targets of miR-199a-3p.

Fig. S7. Interfering with miR-199a-5p function prevents lung fibrosis in vivo.

Fig. S8. Acute and chronic systemic administration of LNA-199a-5p inhibitor does not induce major side effects.

Fig. S9. Raw gel images corresponding to western blot shown on Figure 1.

Fig. S10. Raw gel images corresponding to western blot shown on Figure 3.

Fig. S11. Raw gel images corresponding to western blot shown on Figure 4.

Table S1. List of the TGF-β core signature of 419 coding genes significantly modulated by TGF-β in lung fibroblasts, as identified by Ingenuity Pathway analysis (dataset 1).

Table S2. List of putative lncRNAs significantly modulated by TGF-β in lung fibroblasts (dataset 1).

Table S3. List of the best mature miRNAs significantly modulated by TGF-β in lung fibroblasts following small RNA-seq analysis (Dataset 2).

Table S4. List of themes corresponding to “canonical pathways” annotations associated with TGF-β stimulation of MRC5 lung fibroblasts in presence of a control or DNM3OS-specific gapmer identified by Ingenuity Pathway Analysis.

Table S5. List of themes corresponding to “canonical pathways” annotations identified by Ingenuity Pathway Analysis in response to overexpression of miR-199a-3p, miR-199a-5p or miR-214-3p in human MRC5 pulmonary fibroblasts MRC5.

Table S6. LNA-modified oligonucleotides used for in vivo experiments.

Table S7. Characteristics of 3’UTR psiCHECK-2 plasmid constructions.

Table S8. TaqMan assays used for miRNAs expression analysis.

Table S9. qPCR primers used for gene expression analysis.

## Acknowledgments

The authors gratefully acknowledge the outstanding technical support of V. Magnone, N. Pons, G. Rios (UCA GenomiX platform of the University Cote d’Azur), M. Tardivel (BICeL-Campus HU Facility, Lille), Dr V. Gnemmi and M.H. Gevaert (Pathology Department, Lille), the staffs from the clinical chemistry lab (CHRU Lille) and from the animal care facilities institutions at Sophia Antipolis (IPMC Animal Care Facility) and Lille (High Technology Animal Care Facility, University of Lille 2). The authors thank N. Frandsen (Exiqon) for the design of the different ASOs and for helpful discussion.

## Author contributions

GS, ED, EC, ISH, NM, NP, CC, CVdH, JF performed and analyzed *in vitro* experiments; GS, MB, CLLC performed and analyzed experiments related to lung fibrosis mouse model; GS, ED performed and analyzed experiments related to kidney fibrosis mouse model; ED, GS, NP performed toxicological analysis; NN, KL, BM analyzed transcriptomic datasets and performed biological interpretation; NN and AP performed statistical analysis; MP, BW, LP, FG, CHM, BC, RR, FB, TB, PB, SL, SB gave conceptual advices; NP, BM, CC conceived, designed and coordinated this work and wrote the manuscript.

## References

1. N. Pottier, C. Cauffiez, M. Perrais, P. Barbry, B. Mari, FibromiRs: translating molecular discoveries into new anti-fibrotic drugs., Trends Pharmacol. Sci. 35, 119–26 (2014).

2. T. A. Wynn, Common and unique mechanisms regulate fibrosis in various fibroproliferative diseases., J. Clin. Invest. 117, 524–9 (2007).

3. P. B. Bitterman, C. A. Henke, Fibroproliferative disorders., Chest 99, 81S–84S (1991).

4. S. L. Friedman, D. Sheppard, J. S. Duffield, S. Violette, Therapy for fibrotic diseases: nearing the starting line., Sci. Transl. Med. 5, 167sr1 (2013).

5. M.-L. Bochaton-Piallat, G. Gabbiani, B. Hinz, The myofibroblast in wound healing and fibrosis: answered and unanswered questions., F1000Research 5, 752 (2016).

6. R. T. Kendall, C. A. Feghali-Bostwick, Fibroblasts in fibrosis: novel roles and mediators., Front. Pharmacol. 5, 123 (2014).

7. E. El Agha, A. Moiseenko, V. Kheirollahi, S. De Langhe, S. Crnkovic, G. Kwapiszewska, D. Kosanovic, F. Schwind, R. T. Schermuly, I. Henneke, B. MacKenzie, J. Quantius, S. Herold, A. Ntokou, K. Ahlbrecht, R. E. Morty, A. Günther, W. Seeger, S. Bellusci, Two-Way Conversion between Lipogenic and Myogenic Fibroblastic Phenotypes Marks the Progression and Resolution of Lung Fibrosis, Cell Stem Cell 20, 261–273.e3 (2017).

8. T. A. Wynn, T. R. Ramalingam, Mechanisms of fibrosis: therapeutic translation for fibrotic disease, Nat. Med. 18, 1028–1040 (2012).

9. R. J. Akhurst, A. Hata, Targeting the TGFβ signalling pathway in disease., Nat. Rev. Drug Discov. 11, 790–811 (2012).

10. C. S. King, S. D. Nathan, Idiopathic pulmonary fibrosis: effects and optimal management of comorbidities., Lancet. Respir. Med. 5, 72–84 (2017).

11. G. Hughes, H. Toellner, H. Morris, C. Leonard, N. Chaudhuri, Real World Experiences: Pirfenidone and Nintedanib are Effective and Well Tolerated Treatments for Idiopathic Pulmonary Fibrosis., J. Clin. Med. 5, 78 (2016).

12. S. T. Lehtonen, A. Veijola, H. Karvonen, E. Lappi-Blanco, R. Sormunen, S. Korpela, U. Zagai, M. C. Sköld, R. Kaarteenaho, Pirfenidone and nintedanib modulate properties of fibroblasts and myofibroblasts in idiopathic pulmonary fibrosis., Respir. Res. 17, 14 (2016).

13. E. Conte, E. Gili, E. Fagone, M. Fruciano, M. Iemmolo, C. Vancheri, Effect of pirfenidone on proliferation, TGF-β-induced myofibroblast differentiation and fibrogenic activity of primary human lung fibroblasts., Eur. J. Pharm. Sci. 58, 13–9 (2014).

14. L. Wollin, E. Wex, A. Pautsch, G. Schnapp, K. E. Hostettler, S. Stowasser, M. Kolb, Mode of action of nintedanib in the treatment of idiopathic pulmonary fibrosis, Eur. Respir. J. 45, 1434–1445 (2015).

15. K. V Morris, J. S. Mattick, The rise of regulatory RNA., Nat. Rev. Genet. 15, 423–37 (2014).

16. J. Hausser, M. Zavolan, Identification and consequences of miRNA-target interactions--beyond repression of gene expression., Nat. Rev. Genet. 15, 599–612 (2014).

17. J. T. Mendell, E. N. Olson, MicroRNAs in stress signaling and human disease., Cell 148, 1172–87 (2012).

18. E. van Rooij, S. Kauppinen, Development of microRNA therapeutics is coming of age., EMBO Mol. Med. 6, 851–64 (2014).

19. K. A. Lennox, M. A. Behlke, Cellular localization of long non-coding RNAs affects silencing by RNAi more than by antisense oligonucleotides., Nucleic Acids Res. 44, 863–77 (2016).

20. T. Watanabe, T. Sato, T. Amano, Y. Kawamura, N. Kawamura, H. Kawaguchi, N. Yamashita, H. Kurihara, T. Nakaoka, Dnm3os, a non-coding RNA, is required for normal growth and skeletal development in mice., Dev. Dyn. 237, 3738–48 (2008).

21. C. L. Lino Cardenas, I. S. Henaoui, E. Courcot, C. Roderburg, C. Cauffiez, S. Aubert, M.-C. Copin, B. Wallaert, F. Glowacki, E. Dewaeles, J. Milosevic, J. Maurizio, J. Tedrow, B. Marcet, J.-M. Lo-Guidice, N. Kaminski, P. Barbry, T. Luedde, M. Perrais, B. Mari, N. Pottier, H. S. Scott, Ed. miR-199a-5p Is upregulated during fibrogenic response to tissue injury and mediates TGFbeta-induced lung fibroblast activation by targeting caveolin-1., PLoS Genet. 9, e1003291 (2013).

22. B. B. Moore, C. M. Hogaboam, Murine models of pulmonary fibrosis., Am. J. Physiol. Lung Cell. Mol. Physiol. 294, L152–60 (2008).

23. N. Pottier, T. Maurin, B. Chevalier, M.-P. Puisségur, K. Lebrigand, K. Robbe-Sermesant, T. Bertero, C. L. Lino Cardenas, E. Courcot, G. Rios, S. Fourre, J.-M. Lo-Guidice, B. Marcet, B. Cardinaud, P. Barbry, B. Mari, D.-Y. Jin, Ed. Identification of keratinocyte growth factor as a target of microRNA-155 in lung fibroblasts: implication in epithelial-mesenchymal interactions., PLoS One 4, e6718 (2009).

24. M.-P. Puisségur, N. M. Mazure, T. Bertero, L. Pradelli, S. Grosso, K. Robbe-Sermesant, T. Maurin, K. Lebrigand, B. Cardinaud, V. Hofman, S. Fourre, V. Magnone, J. E. Ricci, J. Pouysségur, P. Gounon, P. Hofman, P. Barbry, B. Mari, miR-210 is overexpressed in late stages of lung cancer and mediates mitochondrial alterations associated with modulation of HIF-1 activity., Cell Death Differ. 18, 465–78 (2011).

25. A. Subramanian, P. Tamayo, V. K. Mootha, S. Mukherjee, B. L. Ebert, M. A. Gillette, A. Paulovich, S. L. Pomeroy, T. R. Golub, E. S. Lander, J. P. Mesirov, Gene set enrichment analysis: a knowledge-based approach for interpreting genome-wide expression profiles., Proc. Natl. Acad. Sci. U. S. A. 102, 15545–50 (2005).

26. A. Greenhough, H. J. M. Smartt, A. E. Moore, H. R. Roberts, A. C. Williams, C. Paraskeva, A. Kaidi, The COX-2/PGE2 pathway: key roles in the hallmarks of cancer and adaptation to the tumour microenvironment., Carcinogenesis 30, 377–86 (2009).

27. A. Akhmetshina, K. Palumbo, C. Dees, C. Bergmann, P. Venalis, P. Zerr, A. Horn, T. Kireva, C. Beyer, J. Zwerina, H. Schneider, A. Sadowski, M.-O. Riener, O. A. MacDougald, O. Distler, G. Schett, J. H. W. Distler, Activation of canonical Wnt signalling is required for TGF-β-mediated fibrosis., Nat. Commun. 3, 735 (2012).

28. T. M. Maher, I. C. Evans, S. E. Bottoms, P. F. Mercer, A. J. Thorley, A. G. Nicholson, G. J. Laurent, T. D. Tetley, R. C. Chambers, R. J. McAnulty, Diminished prostaglandin E2 contributes to the apoptosis paradox in idiopathic pulmonary fibrosis., Am. J. Respir. Crit. Care Med. 182, 73–82 (2010).

29. L. M. Crosby, C. M. Waters, Epithelial repair mechanisms in the lung., Am. J. Physiol. Lung Cell. Mol. Physiol. 298, L715–31 (2010).

30. O. Mungunsukh, R. M. Day, Transforming growth factor-β1 selectively inhibits hepatocyte growth factor expression via a micro-RNA-199-dependent posttranscriptional mechanism., Mol. Biol. Cell 24, 2088–97 (2013).

31. K. A. Bauman, S. H. Wettlaufer, K. Okunishi, K. M. Vannella, J. S. Stoolman, S. K. Huang, A. J. Courey, E. S. White, C. M. Hogaboam, R. H. Simon, G. B. Toews, T. H. Sisson, B. B. Moore, M. Peters-Golden, The antifibrotic effects of plasminogen activation occur via prostaglandin E2 synthesis in humans and mice., J. Clin. Invest. 120, 1950–60 (2010).

32. E. van Rooij, A. L. Purcell, A. A. Levin, Developing microRNA therapeutics., Circ. Res. 110, 496–507 (2012).

33. R. Vittal, J. C. Horowitz, B. B. Moore, H. Zhang, F. J. Martinez, G. B. Toews, T. J. Standiford, V. J. Thannickal, Modulation of prosurvival signaling in fibroblasts by a protein kinase inhibitor protects against fibrotic tissue injury., Am. J. Pathol. 166, 367–75 (2005).

34. Z.-Y. Wu, L. Lu, J. Liang, X.-R. Guo, P. H. Zhang, S.-J. Luo, Keloid microRNA expression analysis and the influence of miR-199a-5p on the proliferation of keloid fibroblasts., Genet. Mol. Res. 13, 2727–38 (2014).

35. E. van Rooij, L. B. Sutherland, J. E. Thatcher, J. M. DiMaio, R. H. Naseem, W. S. Marshall, J. A. Hill, E. N. Olson, Dysregulation of microRNAs after myocardial infarction reveals a role of miR-29 in cardiac fibrosis., Proc. Natl. Acad. Sci. U. S. A. 105, 13027–32 (2008).

36. R. L. Chevalier, M. S. Forbes, B. A. Thornhill, Ureteral obstruction as a model of renal interstitial fibrosis and obstructive nephropathy, Kidney Int. 75, 1145–1152 (2009).

37. S. Hombach, M. Kretz, Non-coding RNAs: Classification, Biology and Functioning., Adv. Exp. Med. Biol. 937, 3–17 (2016).

38. G. Yin, R. Chen, A. B. Alvero, H.-H. Fu, J. Holmberg, C. Glackin, T. Rutherford, G. Mor, TWISTing stemness, inflammation and proliferation of epithelial ovarian cancer cells through MIR199A2/214., Oncogene 29, 3545–53 (2010).

39. H. L. A. Janssen, H. W. Reesink, E. J. Lawitz, S. Zeuzem, M. Rodriguez-Torres, K. Patel, A. J. van der Meer, A. K. Patick, A. Chen, Y. Zhou, R. Persson, B. D. King, S. Kauppinen, A. A. Levin, M. R. Hodges, Treatment of HCV infection by targeting microRNA., N. Engl. J. Med. 368, 1685–94 (2013).

40. X. M. Wang, Y. Zhang, H. P. Kim, Z. Zhou, C. A. Feghali-Bostwick, F. Liu, E. Ifedigbo, X. Xu, T. D. Oury, N. Kaminski, A. M. K. Choi, Caveolin-1: a critical regulator of lung fibrosis in idiopathic pulmonary fibrosis., J. Exp. Med. 203, 2895–906 (2006).

41. M. Drab, P. Verkade, M. Elger, M. Kasper, M. Lohn, B. Lauterbach, J. Menne, C. Lindschau, F. Mende, F. C. Luft, A. Schedl, H. Haller, T. V Kurzchalia, Loss of caveolae, vascular dysfunction, and pulmonary defects in caveolin-1 gene-disrupted mice., Science 293, 2449–52 (2001).

42. M. A. Fernandez-Rojo, G. A. Ramm, Caveolin-1 Function in Liver Physiology and Disease., Trends Mol. Med. 22, 889–904 (2016).

43. E. N. Olson, MicroRNAs as Therapeutic Targets and Biomarkers of Cardiovascular Disease, Sci. Transl. Med. 6, 239ps3–239ps3 (2014).

44. N. Pottier, C. Chupin, V. Defamie, B. Cardinaud, R. Sutherland, G. Rios, F. Gauthier, P. J. Wolters, Y. Berthiaume, P. Barbry, B. Mari, Relationships between early inflammatory response to bleomycin and sensitivity to lung fibrosis: a role for dipeptidyl-peptidase I and tissue inhibitor of metalloproteinase-3?, Am. J. Respir. Crit. Care Med. 176, 1098–107 (2007).

45. K. Le Brigand, K. Robbe-Sermesant, B. Mari, P. Barbry, MiRonTop: mining microRNAs targets across large scale gene expression studies., Bioinformatics 26, 3131–2 (2010).

